# Dominant MLC-causing mutations alter hepaCAM subcellular localization and protein interactome in astrocytes of the developing mouse cortex

**DOI:** 10.1101/2025.08.13.670215

**Authors:** Robert W. Lewis, Breana C. Dogan, Madelyn G. Coble, Amy L. Stanek, Hayli E. Spence-Osorio, Karen L.G. Farizatto, Angie L. Mordant, C. Allie Mills, Laura E. Herring, Katherine T. Baldwin

**Affiliations:** Neuroscience Center, University of North Carolina at Chapel Hill, Chapel Hill, NC 27599; Metabolomics and Proteomics Core Facility, Department of Pharmacology, University of North Carolina at Chapel Hill, Chapel Hill, NC 27599; Department of Cell Biology and Physiology, University of North Carolina at Chapel Hill, Chapel Hill, NC 27599

**Keywords:** astrocyte, development, hepaCAM, leukodystrophy, proteomics

## Abstract

Megalencephalic leukoencephalopathy with subcortical cysts (MLC) is a rare leukodystrophy characterized by early-onset macrocephaly, white matter edema, seizures, and motor and cognitive decline. Missense mutations to hepatic and glial cell adhesion molecule (hepaCAM), also known as GlialCAM, are responsible for approximately twenty-five percent of MLC cases. HepaCAM is highly enriched in astrocytes and plays important roles in astrocyte territory establishment, gap junction coupling, branching organization, synaptic function, and development of the gliovascular unit. The molecular mechanisms through which MLC-causing missense mutations alter hepaCAM function *in vivo* and facilitate MLC pathogenesis during brain development remain largely unknown. Here, we used new viral tools and proximity-based proteomics to examine how three different dominant MLC-causing mutations impact hepaCAM subcellular localization and protein interactome in astrocytes of the developing mouse cortex. We found dramatic defects in hepaCAM distribution throughout the astrocyte, which were common to all mutants tested. We also observed significant changes in protein interactome between wild type and mutant hepaCAM, including decreased association with previously described hepaCAM-interacting proteins Connexin 43 and CLC-2. Moreover, we identified changes in association between hepaCAM and a number of previously undescribed potential hepaCAM-interaction partners, including the epilepsy-associated potassium channel KCNQ2. Collectively, our data provide new insights into hepaCAM function in astrocytes during brain development, reveal altered hepaCAM protein dynamics with MLC missense mutations, and provide a new resource to explore the molecular underpinnings of MLC pathogenesis.

## Introduction

Megalencephalic leukoencephalopathy with subcortical cysts (MLC) is an early-onset and progressive leukodystrophy characterized by macrocephaly, white matter edema, seizures, and motor and cognitive decline^1,2^. A majority of MLC cases are caused by homozygous recessive mutations in MLC1, an astrocyte-specific transmembrane protein of unknown function^3^. Approximately one quarter of MLC cases are caused by mutations in hepatic and glial cell adhesion molecular (hepaCAM)^1,4,5^, an astrocyte-enriched cell adhesion molecule with important roles in astrocyte territory establishment, gap junction coupling, branching organization, synaptic function, neurite outgrowth, and development of the gliovascular unit^6–8^. At the cellular level, MLC appears to be a disorder of astrocyte ion and water regulation^9^. A growing body of evidence supports this notion, including clinical findings^1,9,10^, and phenotypes of MLC mouse models^11–14^. Understanding the molecular basis of this dysfunction is hindered by an incomplete understanding of how MLC-causing mutations impact MLC1 and hepaCAM protein function in astrocytes during brain development.

HepaCAM is a single pass transmembrane protein and member of the immunoglobulin super family. In the mouse cortex, hepaCAM is strongly enriched in astrocytes and is expressed throughout the membrane, including major branches, leaflets, endfeet, and cell-cell junctions^6,7^. Interestingly, hepaCAM is required for proper expression and localization of MLC1 both *in vitro* and *in vivo*^11,15^. An *in vitro* study using HeLa cells and U251N, a tumor cell line, found that hepaCAM functions as a chaperone-like protein for MLC1, stabilizing its expression both in the endoplasmic reticulum (ER) and at the membrane^16^. In addition to its regulation of MLC1, hepaCAM is required for proper localization of a growing list of transmembrane proteins that play important roles in ion and fluid homeostasis, including the chloride channel CLC-2^17,18^, and gap junction protein Connexin 43 (Cx43)^6,19^. During normal brain development, hepaCAM is required for astrocyte territory establishment and proper gap junction coupling^6^. HepaCAM also functions at synapses to regulate synaptic strength^6^ and on astrocyte-secreted exosomes to promote axonal outgrowth^8^.

Most MLC-causing mutations to hepaCAM are missense mutations located in the extracellular IgV domain^20–22^. These mutations may be dominant or recessive depending on the location and specific amino acid substitution. Homozygous recessive mutations to hepaCAM cause MLC Type 2a, a progressive disease similar to MLC Type 1, which is caused by mutations to MLC1^23^. In contrast, heterozygous dominant mutations to hepaCAM cause MLC Type 2b, a milder form of the disease with a remitting phenotype and more frequent comorbidity with autism spectrum disorder^1,5,24^. A strong body of i*n vitro* studies using HeLa cells or primary mouse astrocytes have shown that hepaCAM is enriched at cell-cell junctions, where it typically interacts with itself both in *cis* and in *trans*, and demonstrated that MLC-causing hepaCAM mutations impair the ability of hepaCAM to localize to these cell-cell junctions^5,21,22,25^. Recessive hepaCAM mutations, and some dominant mutations are thought to impair *cis* interaction based on their location within the IgV domain, while other dominant mutations impair *trans* interaction^20^. *In vivo*, mice that express the dominant G89S mutation showed altered hepaCAM localization in the molecular layer of the cerebellum^18^. To date, how MLC-causing hepaCAM mutations impact protein localization and function in astrocytes *in vivo*, particularly within the context of brain development, remains unexplored.

Here, we investigated the impact of dominant MLC-causing missense mutations on hepaCAM protein dynamics in astrocytes of the developing mouse cortex. Our findings reveal altered subcellular localization of mutant hepaCAM in astrocytes and significant changes in association with key membrane proteins. Through proximity-based quantitative proteomics, we describe how the hepaCAM interactome changes across mutations and provide a valuable resource for exploring the molecular basis of MLC pathogenesis.

## Results

### MLC-causing missense mutations alter hepaCAM localization in astrocytes co-cultured with neurons

We previously investigated the function of hepaCAM in astrocyte development using an astrocyte-neuron co-culture system and exogenous expression of hepaCAM mutants^6^. Using this system, we observed substantially altered subcellular localization of both dominant G89S and recessive R29Q mutant versions of human hepaCAM protein. Both mutants showed strong, relatively homogenous expression throughout the cells, in stark contrast to the punctate expression of endogenous hepaCAM or exogenously expressed wild type (WT) hepaCAM^6^. Both G89S and R92Q impair hepaCAM *cis* interaction^20^. To test whether mutations that impair hepaCAM *trans* interaction behave similarly, we performed site-directed mutagenesis to introduce different dominant MLC-causing mutations into vectors that expressed human hepaCAM-HA under control of the gfaABC1D promoter (**Figure 1A**). Both Q56P and D128N impair *trans* interaction, and D128N introduces a new N-glycosylation site^20^. We expressed these *trans*-mutants in astrocytes co-cultured with neurons and examined their localization in comparison to WT and G89S hepaCAM-HA using HA immunolabeling (**Figure 1A-B**). We also co-transfected a plasmid expressing green fluorescent protein (GFP) to visualize cellular structure.

**Figure 1:**
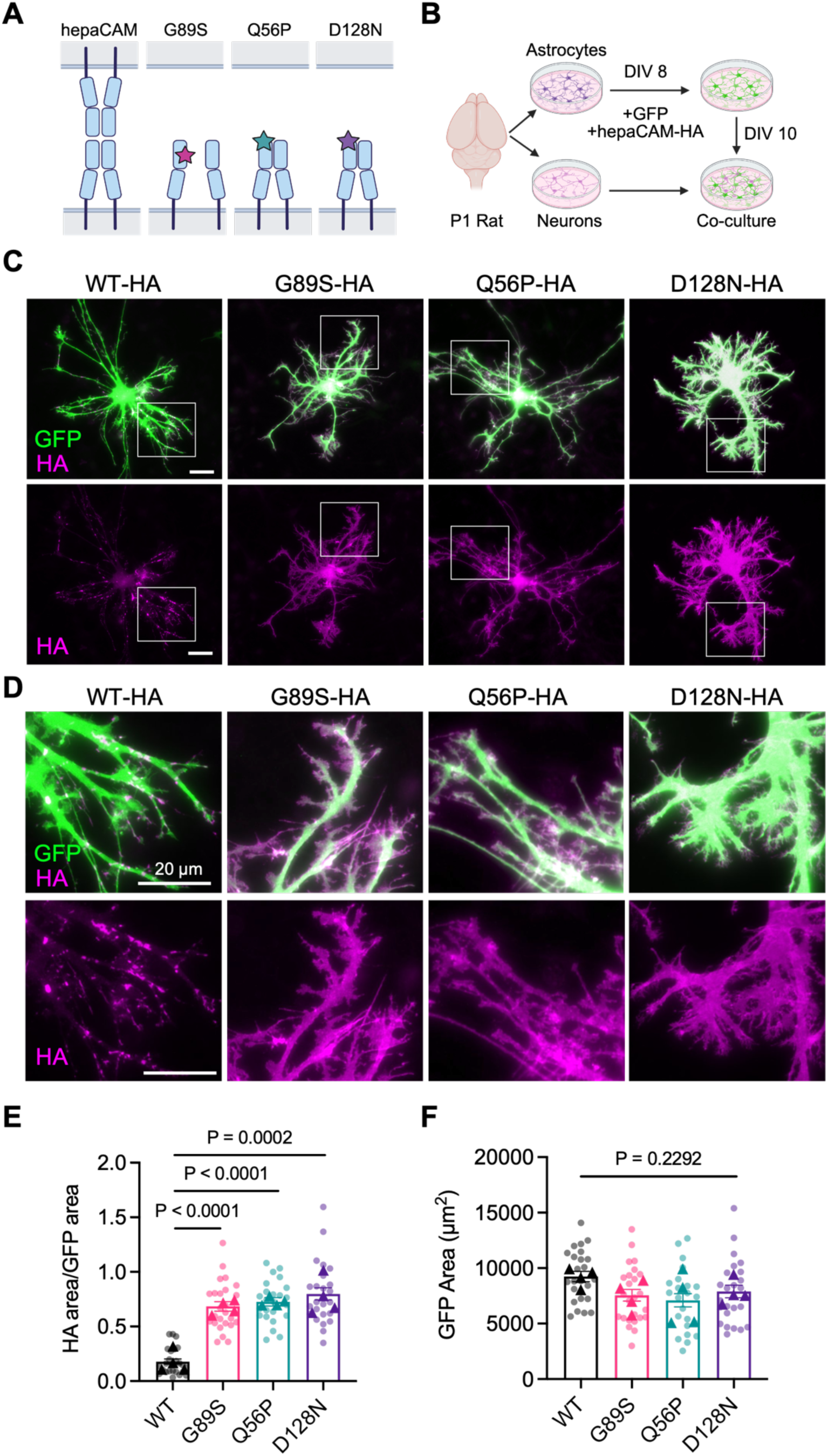
MLC-causing mutations alter hepaCAM localization in astrocytes co-cultured with neurons. A) Diagram showing wild-type (WT) and mutant hepaCAM interactions at astrocyte-astrocyte contacts. WT hepaCAM interacts with itself in *cis* and *trans*. G89S mutation impairs *cis* interaction, which may impair *trans* interaction. Q56P and D128N mutations impair *trans* interaction. B) Co-culture workflow. HA tagged hepaCAM constructs are transfected into astrocytes after 8 days in vitro (DIV 8) prior to co-culture with neurons (DIV 10). C) Representative images of astrocytes transfected with GFP (green) and HA-tagged hepaCAM constructs (magenta) and co-cultured with neurons. Scale bar 20 µm. The region within the white box for each image is enlarged in panel D). E-F) Quantification of E) fraction of GFP area occupied by HA signal and F) GFP cell area N = 4 experiments, 5-8 cells/condition/experiment. One-way ANOVA with Tukey’s HSD.

While WT hepaCAM showed a characteristic punctate distribution throughout the branches, G89S, Q56P and D128N mutants were broadly and homogenously distributed throughout the cell (**Figure 1C**). At the tips of branches, mutant hepaCAM was visible in lamellipodia-like protrusions that exceeded the territory of the cytosolic GFP signal (**Figure 1D**). Analysis of HA signal area normalized to cell (GFP) area revealed a substantial and significant increase in hepaCAM signal area in all three mutants (**Figure 1E**), with no significant change in cell area (**Figure 1F**). Thus, in astrocytes co-cultured with neurons, dominant MLC-causing mutations that impair either *cis* or *trans* homophilic hepaCAM interaction similarly alter hepaCAM localization.

### Dominant MLC-causing missense mutations alter hepaCAM protein distribution in the developing mouse cortex

To understand how MLC-causing mutations to hepaCAM impact hepaCAM localization and function during brain development, we transitioned our studies to an *in vivo* model. We chose the developing mouse visual cortex as we have previously characterized the expression and function of hepaCAM in this brain region during postnatal development^6^. To express WT and mutant human hepaCAM in astrocytes of the developing mouse brain, we packaged hepaCAM-expressing plasmids into PHP.eB serotype adeno-associated virus (AAV) under control of the gfaABC1D promoter. We also fused TurboID to the C-terminus to enable proximity labeling in the presence of biotin, followed by an HA tag to detect exogenous protein expression (**Figure 2A**). Due to the nature of our viral overexpression approach, we focused our studies on dominant, rather than recessive mutations. We performed bilateral intracortical injection of hepaCAM-expressing AAVs at postnatal day 1 (P1) and collected brains for immunohistochemistry at P21, a juvenile timepoint when synaptogenesis and astrocyte maturation are largely complete and roughly corresponding to one year of age in terms of human brain development ^26,27^. For astrocyte membrane visualization, brains were co-transduced with mCherry-CAAX-expressing AAVs (**Figure 2B**).

**Figure 2:**
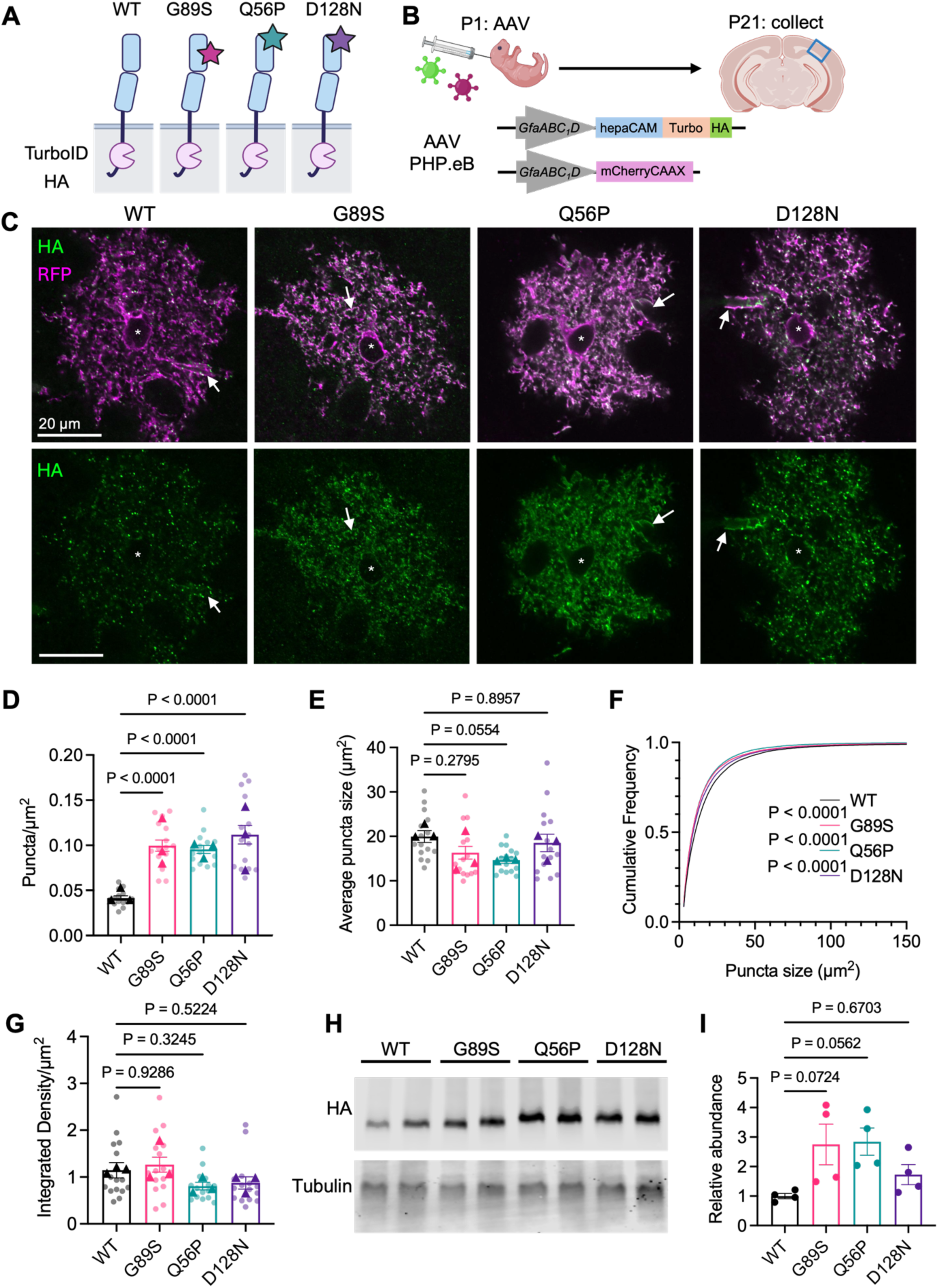
Dominant MLC-causing mutations alter hepaCAM localization *in vivo*. A) Schematic of hepaCAM-TurboID-HA fusion proteins used for *in vivo* expression. B) P1 mice were administered intracortical AAV to express hepaCAM-Turbo-HA fusion proteins and mCherryCAAX. Brains were collected at P21 and astrocytes in the mouse visual cortex (VCX; blue box) analyzed. C) Representative maximum-projection images (3-slices, 0.34 µm step size) of virally-transduced astrocytes in layer 5 of the mouse VCX at P21 with mCherryCAAX in magenta and HA in green. Asterisk indicates astrocyte cell body and arrows indicate astrocyte endfeet. Scale bar 20 µm. D) Analysis of HA puncta density and E) average puncta area for WT and mutant hepaCAM. F) Cumulative frequency distribution of HA puncta size. G) Integrated density normalized to astrocyte area. For D-F, n = 3 animals/condition (triangles), with 5 cells/animal (dots). One-way ANOVA with Tukey’s HSD. H) Western blot of P21 cortical lysates, showing expression of HA and beta-Tubulin. I) Quantification of HA intensity, normalized to beta-tubulin and presented as signal abundance relative to WT. n = 4 animals per condition.

Endogenous mouse hepaCAM shows a punctate pattern of expression throughout the astrocyte membrane (**Figure S1A-B**), including at astrocyte endfeet (**Figure S1C**)^6^. WT hepaCAM-Turbo-HA (hereafter referred to as WT) showed a similar expression pattern (**Figure 2C**) and was strongly associated with endogenous hepaCAM puncta (**Figure S1B-C**), suggesting that the addition of C-terminal Turbo-HA and exogenous viral expression does not disrupt hepaCAM localization. While the G89S, Q56P, and D128N mutants all displayed a punctate expression pattern, the puncta density was significantly increased for all three mutants compared to WT (**Figure 2D**). Though the average puncta size was unchanged (**Figure 2E**), the puncta size cumulative frequency distribution for all three mutants was significantly left-shifted, indicating a greater proportion of smaller-sized puncta compared to WT (**Figure 2F**). The overall signal intensity for each mutant, normalized to cell area, was unchanged (**Figure 2G**). Western blot analysis of whole cortical lysates showed a trend towards increased expression of the G89S and Q56P mutants, but this did not reach statistical significance (**Figure 2H-I**). Interestingly, both the Q56P and D128N mutants showed an upward band shift, which could be indicative of changes to protein glycosylation or other post-translational modifications. Collectively, these results demonstrate similar changes in hepaCAM protein distribution *in vivo* amongst all three dominant mutations tested.

### Decreased co-localization of mutant hepaCAM with Connexin

HepaCAM expression is required for the proper localization of a growing list of transmembrane proteins *in vivo*^28^, including the astrocyte gap junction protein Connexin 43 (Cx43)^6^. Frequent colocalization of hepaCAM and Cx43 is observed in visual cortex astrocytes via confocal and super resolution microscopy, and deletion of hepaCAM from astrocytes alters subcellular Cx43 localization^6^. *In vitro*, R92Q and R92W hepaCAM mutants impair the ability of hepaCAM to localize to cellular junctions^19^. To determine whether G89S, Q56P, or D128N mutations impact hepaCAM association with Cx43 association *in vivo*, we co-labeled tissue sections with HA and Cx43 and assayed the degree of co-localization in transduced astrocytes in Layer V of the mouse visual cortex using Synbot^29^ (**Figure 3A**). Consistent with previous findings, we observed frequent co-localization between WT hepaCAM and Cx43 (**Figure 3B**). For all three mutants tested, we found a significant reduction in the percentage of HA puncta that co-localized with Cx43 (**Figure 3B**). Interestingly, there was no change in the overall density of Cx43 puncta in astrocytes expressing mutant hepaCAM constructs (**Figure 3C**). Q56P-expressing astrocytes showed a significant decrease in the density of co-localized HA and Cx43 puncta, but no significant difference was observed for G89S and D128N (**Figure 3D**). Whether the decrease in percentage of hepaCAM puncta co-localized with Cx43 is driven by a decreased ability of mutant hepaCAM to interact with Cx43, or by a redistribution of hepaCAM protein upstream of Cx43 interaction is unclear. Because confocal microscopy lacks the resolution to detect protein-protein interaction, these results do not provide information on whether MLC-causing mutants impact direct interaction between hepaCAM and Cx43. Quantitative biochemical data is needed to answer this question.

**Figure 3:**
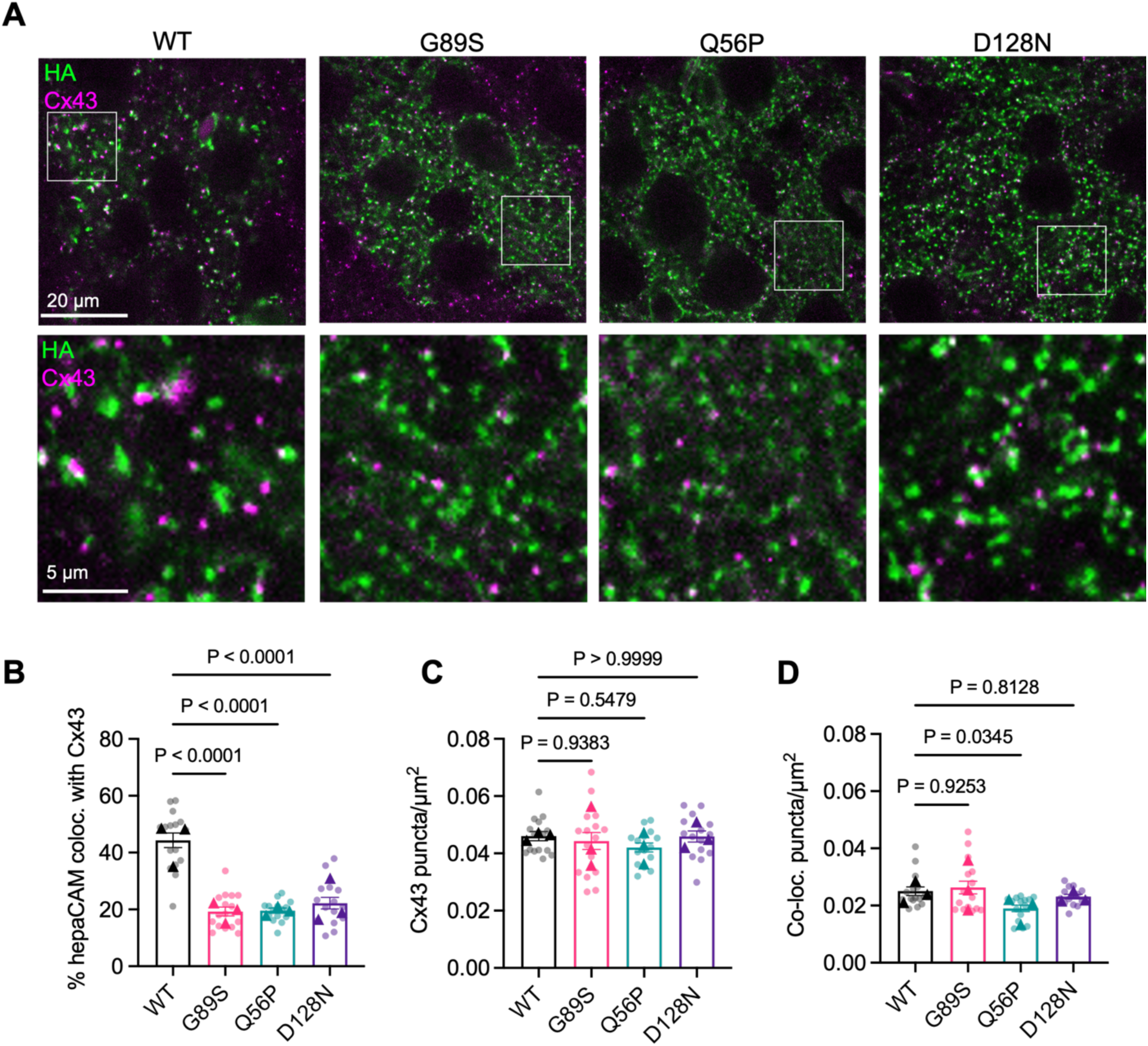
MLC-causing mutations show decreased co-localization with Connexin 43 *in vivo*. A) Representative maximum-projection images (3-slices, 0.34 µm step size) of transduced astrocytes in layer 5 of the mouse VCX at P21, with HA in green and Cx43 in magenta. Scale bar 20 µm. B) Percentage of hepaCAM puncta co-localized with Cx43 puncta. C) Cx43 puncta density. D) Density of co-localized puncta. n = 3 animals/condition (triangles), with 5 cells/animal (dots). One-way ANOVA with Tukey’s HSD.

### Unbiased proximity-based proteomic investigation of hepaCAM and hepaCAM mutant protein interactome

To better determine how MLC-causing mutations impact hepaCAM protein dynamics *in vivo*, we transitioned to an unbiased approach using quantitative proximity-based proteomics with astrocyte-targeted TurboID. In the presence of ATP and biotin, TurboID enables protein labeling within a small (∼10nm) radius through biotinylation^30,31^. Subsequent precipitation with streptavidin-coated beads enables quantitative proteomic analysis (**Figure S2A**).

To perform proximity labeling with WT and mutant hepaCAM probes in astrocytes during brain development, we injected P1 mouse pups intracortically with AAVs to express WT, G89S, Q56P, D128N, or a cytosolic Turbo control under the control of the human minimal GFAP promoter (**Figure 4A**). At P18, mice received daily subcutaneous injection of biotin (24 mg/kg) for three days with brains collected for analysis at P21 (**Figure 4B**). This protocol induced robust biotinylation of astrocytes throughout the cortex, as visualized by fluorophore-conjugated streptavidin (**Figure 4C**). At the cellular level, Turbo-HA was expressed broadly throughout the cytosol, including the cell soma and branches, with similarly localized streptavidin labeling (**Figure 4D**). Consistent with our observations in Figure 2, WT-HA was expressed in a punctate pattern throughout the cell arbor, with little expression in the soma (**Figure 4D**). The streptavidin labeling, which occurred over three days, covered a larger domain than the HA signal, which detects only the current hepaCAM location at time of tissue fixation. This suggests that hepaCAM is a dynamic protein with a broad impact throughout the membrane. All three hepaCAM mutant proteins demonstrated similarly effective biotin labeling (**Figure 4D**).

**Figure 4:**
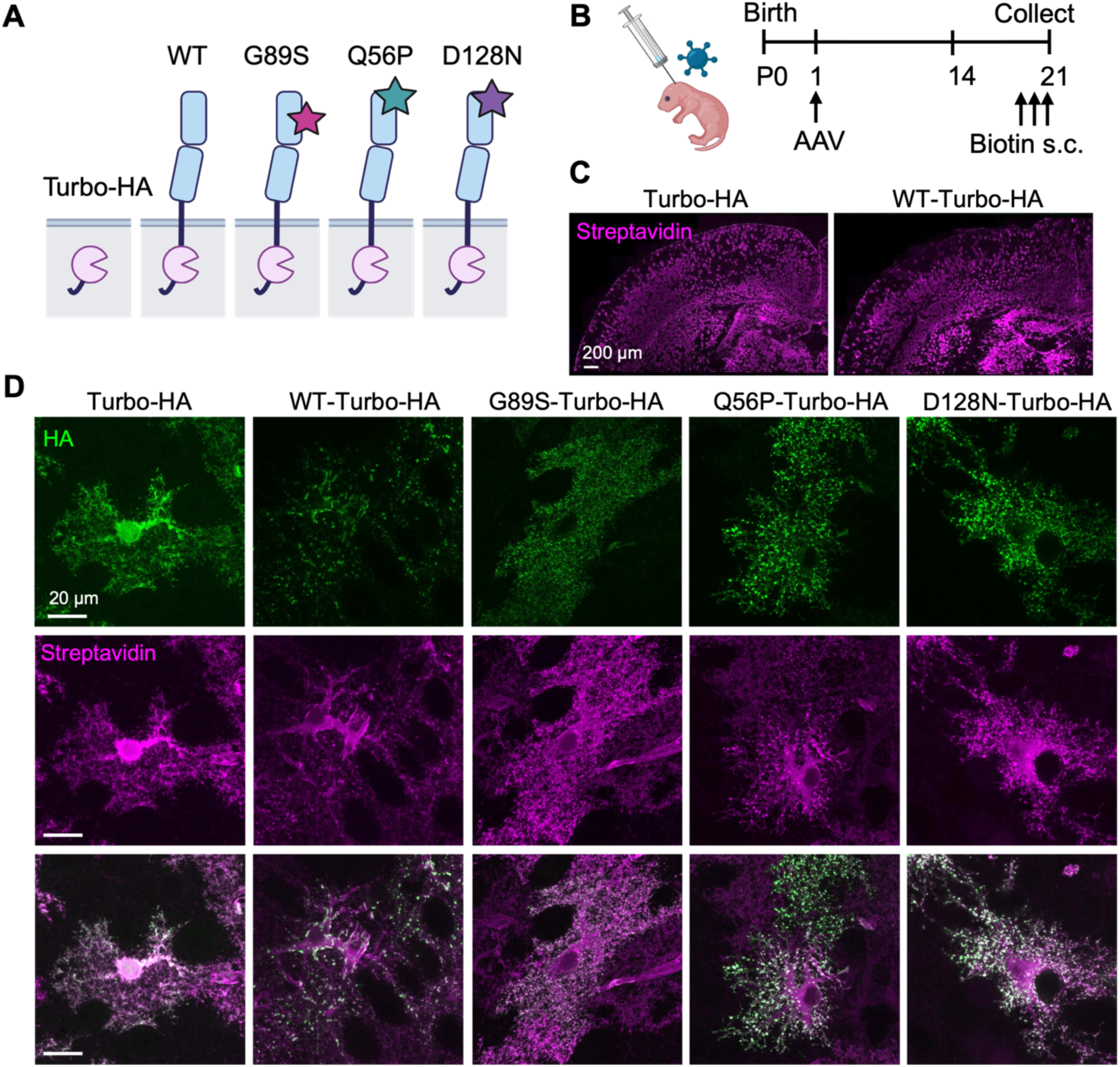
Tool validation for hepaCAM TurboID. A) Diagram of the 5 different conditions used in all TurboID experiments. A cytosolic TurboID-HA driven by the gfaABC1D promoter is used as a control. WT and mutant constructs are identical to figure 2A. B) Timeline of AAV injection and subcutaneous (s.c.) biotin injection. C) Tile scan confocal images of the mouse cortex from Turbo-HA and WT-Turbo-HA transduced brains following biotin administration and labelling with fluorophore-conjugated streptavidin (magenta). Scale bar 200 µm. D) Representative images of astrocytes in the mouse visual cortex following AAV injection and biotin administration. HA in green and streptavidin in magenta. Scale bar 20 µm.

To identify and quantify biotinylated proteins, we dissected the cortex (n = 4 mice/condition) and performed streptavidin pulldowns to isolate biotinylated proteins for liquid chromatography tandem mass spectrometry (LC-MS/MS) (**Figure S2B**). After removing known contaminants and filtering out proteins with only one unique peptide hit, we identified a total of 3720 unique proteins across conditions (**Table S1**). We used log2-transformed Label-free quantity (LFQ) intensities for each individual sample to calculate the log2 fold change (log_2_FC) for WT and mutant hepaCAM conditions compared to cytosolic TurboID. In the WT condition, we identified 1,205 proteins that were significantly enriched over TurboID (p < 0.05, log_2_FC > 1) (**Figure S2C**). We compared our results to a study that used hepaCAM co-immunoprecipitation from adult whole brain lysates to characterize the hepaCAM interactome^28^. Of the 21 proteins described in this study, we identify 19/21 of these proteins in our dataset. Of these, 14/19 were significantly enriched over TurboID, including MLC1, CLC-2 (Clcn2), Cx43 (Gja1), GPRC5B, and GPR37L1 (**Table 1**). One of the proteins that we did not detect, GPR37, is expressed only by oligodendrocytes^32,33^, providing confirmation of the specificity of our approach for targeting astrocytes. The cell adhesion molecule CD44 topped the list as the most enriched protein (log_2_FC = 5.51, p = 0.00035). In cultured astrocytes, CD44 has been shown to regulate astrocyte morphology via Rac1^34^, but an interaction between hepaCAM and CD44 has not been described. To validate our proteomic findings and determine whether CD44 closely associates with endogenous hepaCAM *in vivo*, we performed three-color stimulated emission depletion (STED) in the visual cortex of Aldh1L1eGFP mice at P21. We observed co-localization of hepaCAM and CD44 at super resolution, with hepaCAM signal organized around CD44 in the stereotypical “cupping” fashion that we previously observed with Cx43^6^ (**Figure S3**). CD44 was also the top protein hit in G89S and Q56P compared to TurboID and placed second in D128N behind Drp2 (**Figure S2D-F, Table S1**).

**Table 1:**
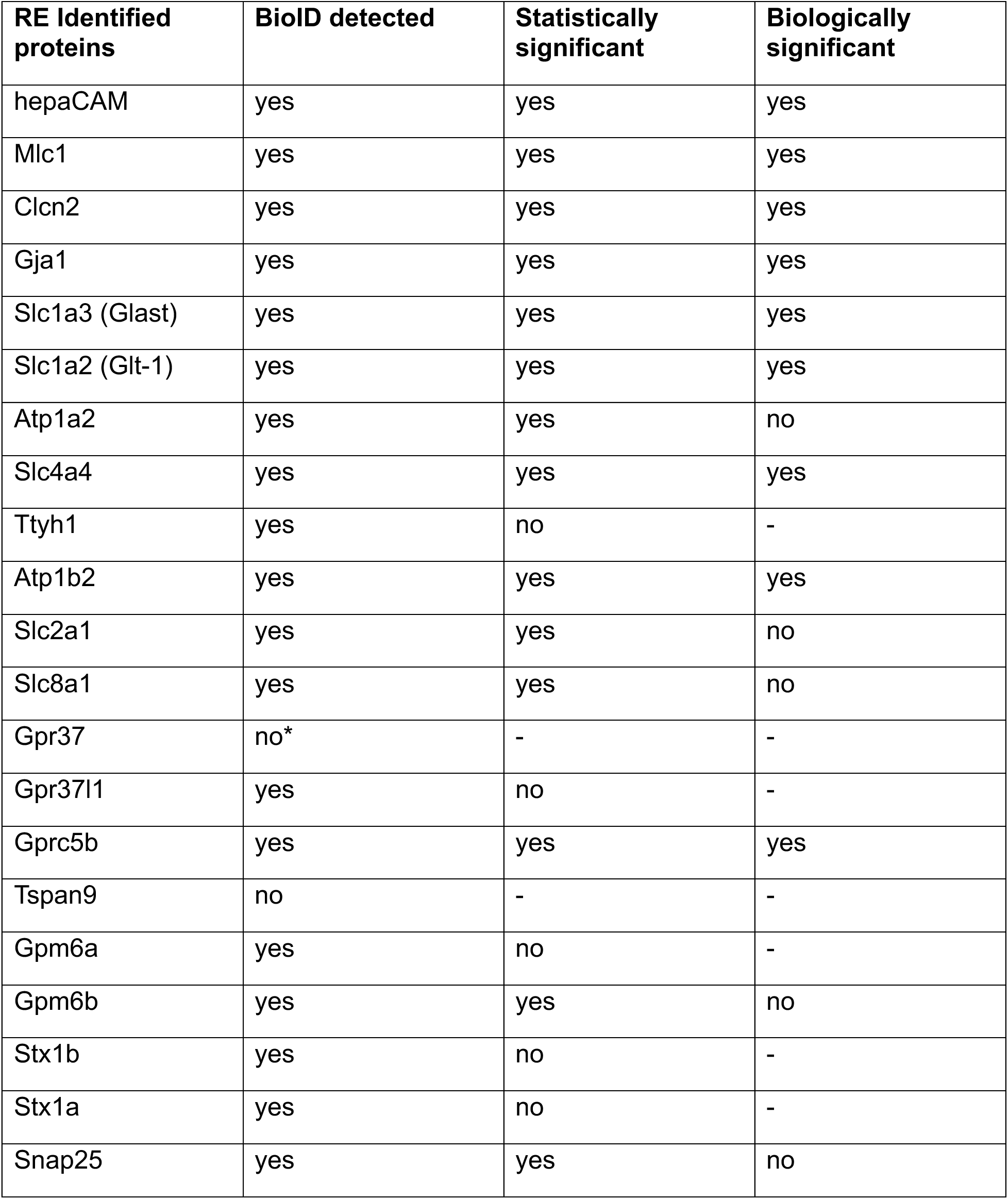
Comparison of BioID results with previously published hepaCAM interactome. *Gpr37 is an oligodendrocyte-specific protein that is not expressed in astrocytes.

### Dominant MLC-causing mutations alter hepaCAM protein interactome

Next, to determine how MLC-causing mutations impact hepaCAM protein association, we compared target protein abundance between WT and hepaCAM mutant conditions (**Figure 5A-C**). For each comparison, we considered proteins with a log_2_FC > 0.585 in the mutant condition compared to WT (corresponding to 50% increase over WT) and p-value < 0.05 to be significantly increased in abundance. We considered proteins with a log_2_FC < -0.585 and p-value < 0.05 to be significantly decreased in abundance. For all comparisons, we excluded proteins that were not significantly enriched above TurboID in at least one condition. Of the 3720 proteins in our dataset, only a small fraction of these were significantly altered between WT and mutant conditions. Q56P had the highest number of increased proteins (66, 1.8%), followed by G89S (49, 1.3%), and D128N (40, 1.1%) (**Figure 5A-C**). There were 16 proteins with significantly increased abundance in all three mutant conditions, including CD44, and known-hepaCAM interacting proteins GPRC5B, TTYH1, and GPR37L1^4,28^ (**Figure 5D-E**). Slightly more proteins showed decreased abundance in mutant conditions with D128N having the highest (97, 2.6%), followed by G89S (85, 2.2%), and Q56P (75, 2.0%) (**Figure 5A-C**). There were 49 proteins with significantly decreased abundance common to all mutant conditions, including known hepaCAM-interacting proteins CLC-2 and Cx43 (**Figure 5D-E**). We did not observe any significant change in MLC1 abundance across WT and mutant protein interactomes (**Figure 5E**).

**Figure 5:**
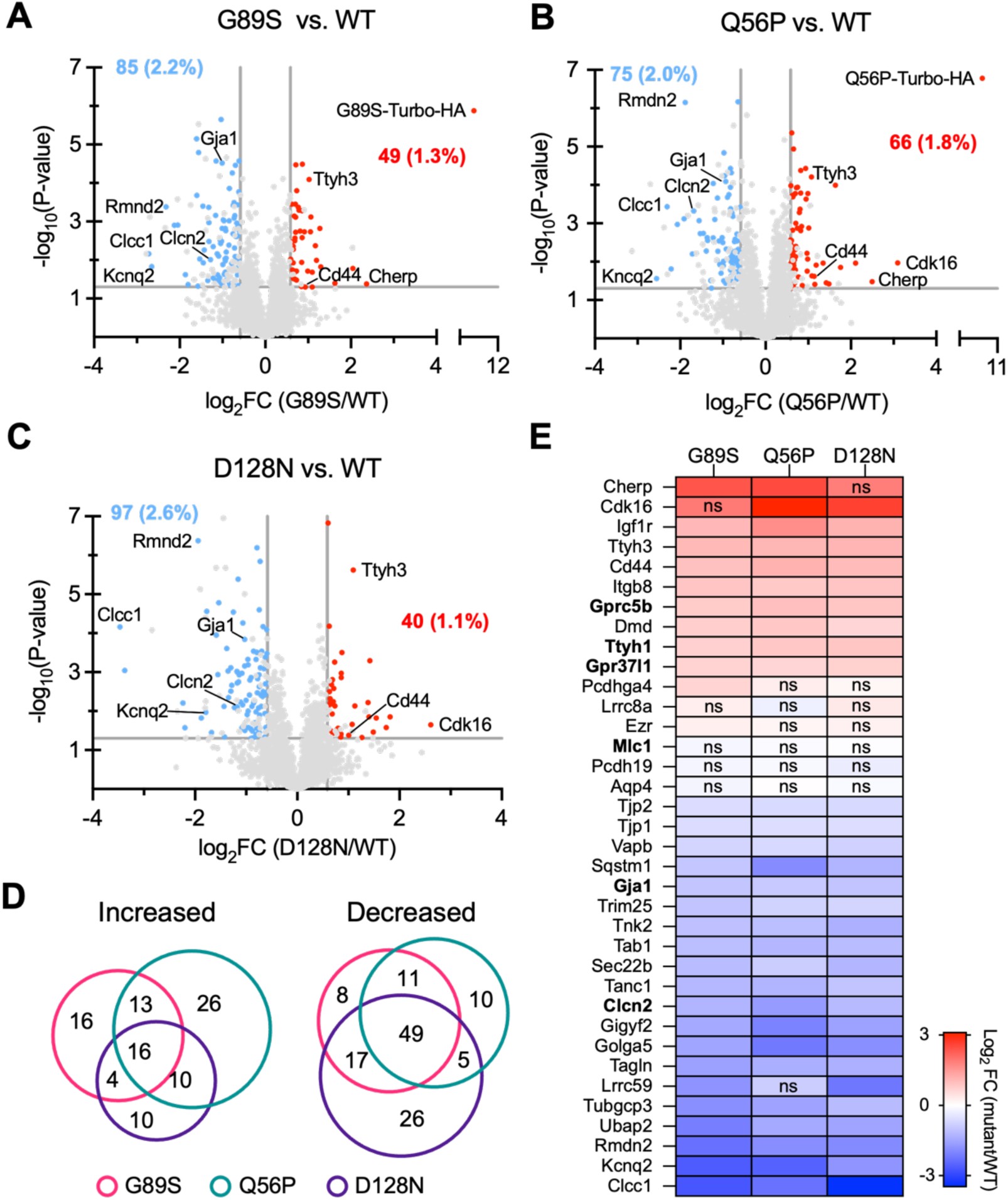
MLC-causing mutations alter hepaCAM protein interactome in vivo. A-C) Volcano plots comparing differentially enriched proteins between WT and A) G89S, B) Q56P, and C) D128N TurboID probes. Red dots indicate proteins significantly enriched in the mutant conditions (p < 0.05, log_2_FC > 0.585). Blue dots indicate proteins significantly reduced in the mutant conditions (p < 0.05, Log_2_FC < -0.585). Any proteins that were not significantly enriched in either the WT or mutant conditions compared to the cytosolic TurboID control were excluded (gray dots that appear in red and blue sections of graph). D) Venn diagram showing number of overlapping and unique proteins that were increased or decreased for each WT-mutant comparison. E) Heat map comparing log_2_FC of increased, decreased, and unchanged protein targets in each WT vs. mutant comparison. ns = p > 0.05.

One of the most consistently decreased proteins across all mutant conditions was the voltage-gated potassium channel KCNQ2 (**Figure 5E**). Mutations in KCNQ2 cause developmental epileptic encephalopathy (DEE)^35,36^, which is of interest given the prevalence of seizures in MLC patients. While KCNQ2 function has been primarily studied in neurons, a recent study described a role for glial KCNQ channels in controlling neuronal output^37^. Thus far, no interaction between hepaCAM and KCNQ2 or any other potassium channel has been described. To validate our proteomic findings, we first performed IHC of virus-injected brains to visualize exogenous WT or mutant hepaCAM (HA) and endogenous KCNQ2 (**Figure 6A**). In astrocytes transduced with WT hepaCAM, we detected frequent co-localization between HA and KCNQ2 signals (**Figure 6B**). In contrast, all three mutants showed significantly reduced co-localization with KCNQ2 (**Figure 6B**), consistent with our proteomic findings. To determine whether KCNQ2 closely associates with endogenous hepaCAM in astrocytes, we performed three-color STED in the visual cortex of Aldh1L1eGFP mice at P21 to detect GFP, hepaCAM, and KCNQ2. Consistent with other studies, KCNQ2 was abundantly expressed in neurons. We also detected strong KCNQ2 signal within astrocyte branches (**Figure 6C**, large inset). Astrocytic KCNQ2 showed regular co-localization with hepaCAM at the level of super resolution. Like with CD44 (**Figure S3**) and Cx43^6^, hepaCAM was organized around larger KCNQ2 puncta in a distinct “cupping” pattern (**Figure 6C**, small insets). Collectively these results identify KCNQ2 as a potential interaction partner for hepaCAM and demonstrate the utility of our dataset for uncovering new biology related to hepaCAM and MLC.

**Figure 6:**
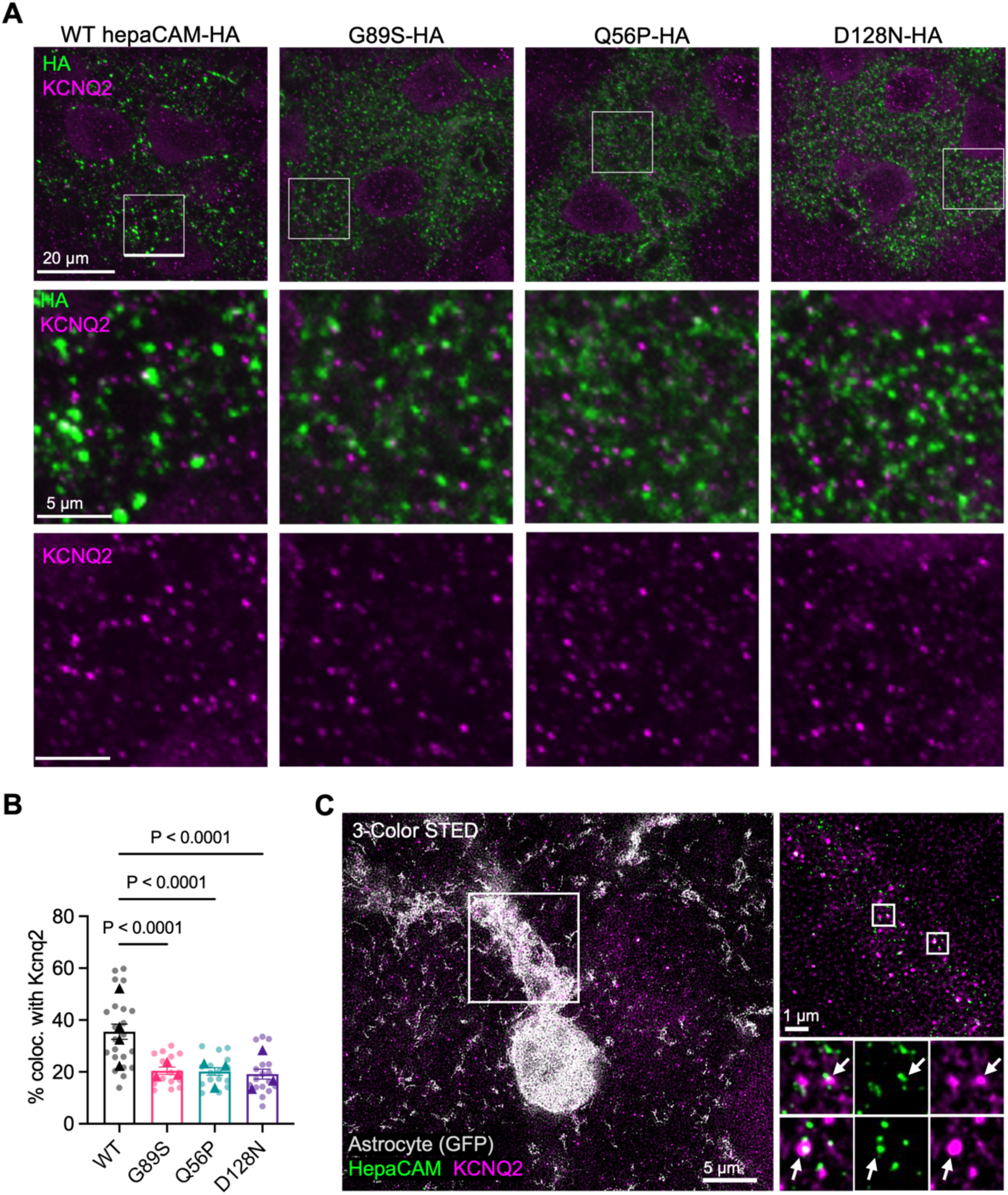
MLC-causing mutations decrease hepaCAM-KCNQ2 colocalization *in vivo*. A) Representative images of single optical sections of virally-transduced astrocytes in layer 5 of the mouse VCX at P21 with HA in green and KCNQ2 in magenta. B) Percentage of hepaCAM puncta co-localized with KCNQ2 puncta. n = 3 animals/condition (triangles), with 5 cells/animal (dots). One-way ANOVA with Tukey’s HSD. C) Representative three-color STED images of astrocytes L5 visual cortex from Aldh1L1eGFP mice. GFP-labeled astrocyte (gray), endogenous hepaCAM (green), KCNQ2 (magenta).

### Analysis of increased and decreased protein networks in hepaCAM mutants

To gain additional insight into our proteomic findings, we performed network analysis of significantly increased (**Figure 7**) and significantly decreased (**Figure 8**) proteins. For analysis of increased proteins, we included all proteins that were significantly increased in at least one mutant condition and visualized these as a single network in Cytoscape with hepaCAM-Turbo-HA as a central node. We performed STRING analysis to identify known and predicted interactions between target nodes (**Figure 7A-B**). This network contained significantly more interactions than expected by chance (protein-protein interaction (PPI) p-value = 7.77e-16), indicating that many of the proteins in the group are biologically related. The top two biological processes identified by Gene Ontology (GO) analysis were cell-cell adhesion and cell migration, (**Figure 7C**) which we visualized within the network by coloring the nodes associated with each term (**Figure 7D**). We also found enrichment of the cellular component GO term Lamellipodium (FDR = 1.8e-4), consistent with our *in vitro* observations of mutant subcellular localization in lamellipodia-like structures (**Figure 1C**). The full list of enriched GO terms in all categories is available in **Table S2**. These results suggest that MLC-causing mutations increase association between hepaCAM and other proteins involved in cell adhesion and migration, though the impact that this has on the function of associated proteins remains to be determined.

**Figure 7:**
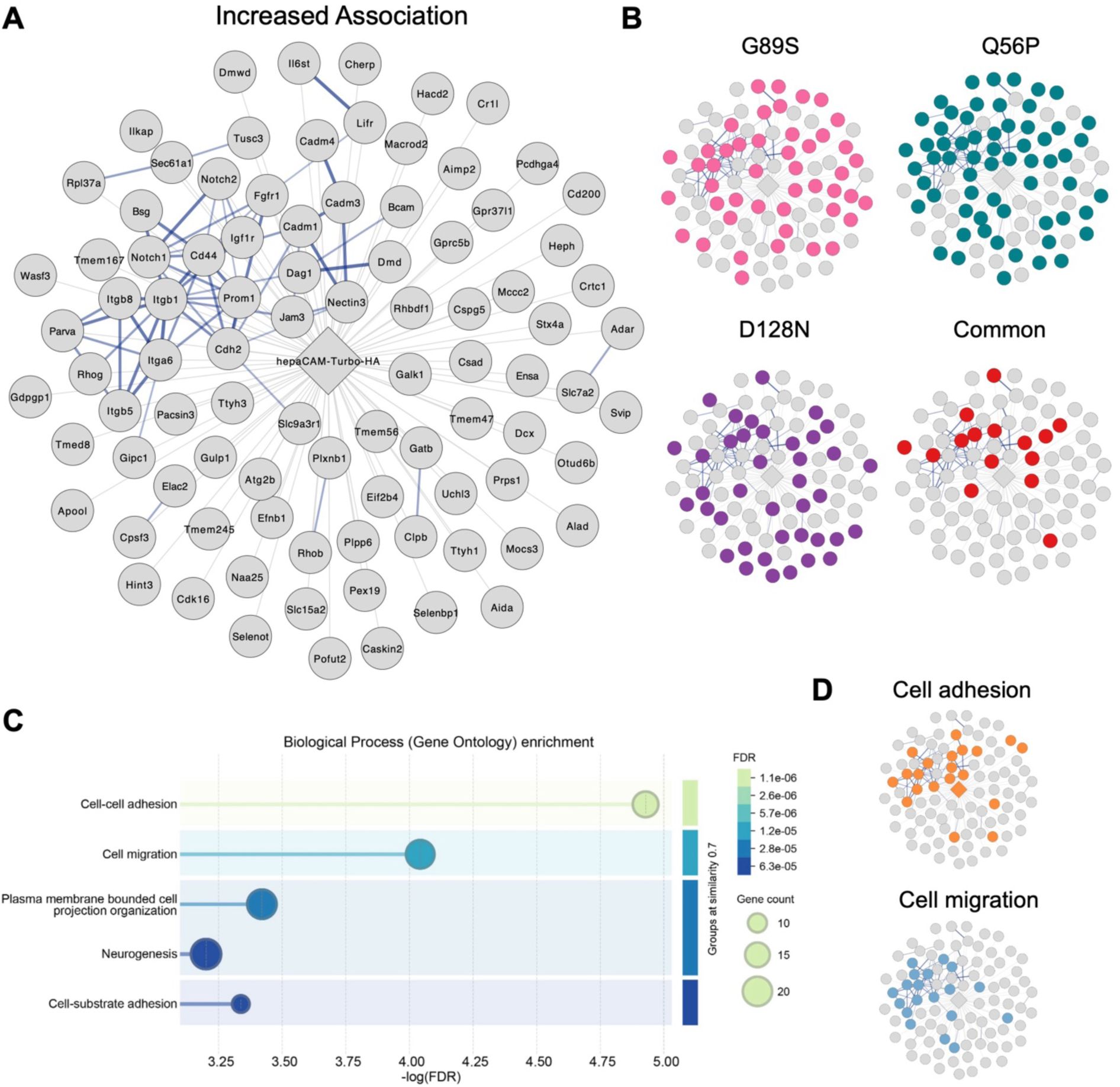
Network analysis reveals hepaCAM mutants preferentially increase interactions with proteins implicated in cell adhesion and migration. A) Significantly increased proteins from the three mutant conditions vs. WT depicted as an interaction network with hepaCAM-Turbo-HA as the central node. Blue lines represent STRING analysis to detect predicted protein-protein interactions. B) Location of individual mutant vs. WT proteins within the network and proteins common to all three mutant vs. WT networks. C) Gene Ontology analysis of Biological Process enrichment based on -log(FDR). D) Location of proteins associated with top 2 GO terms within the network.

**Figure 8:**
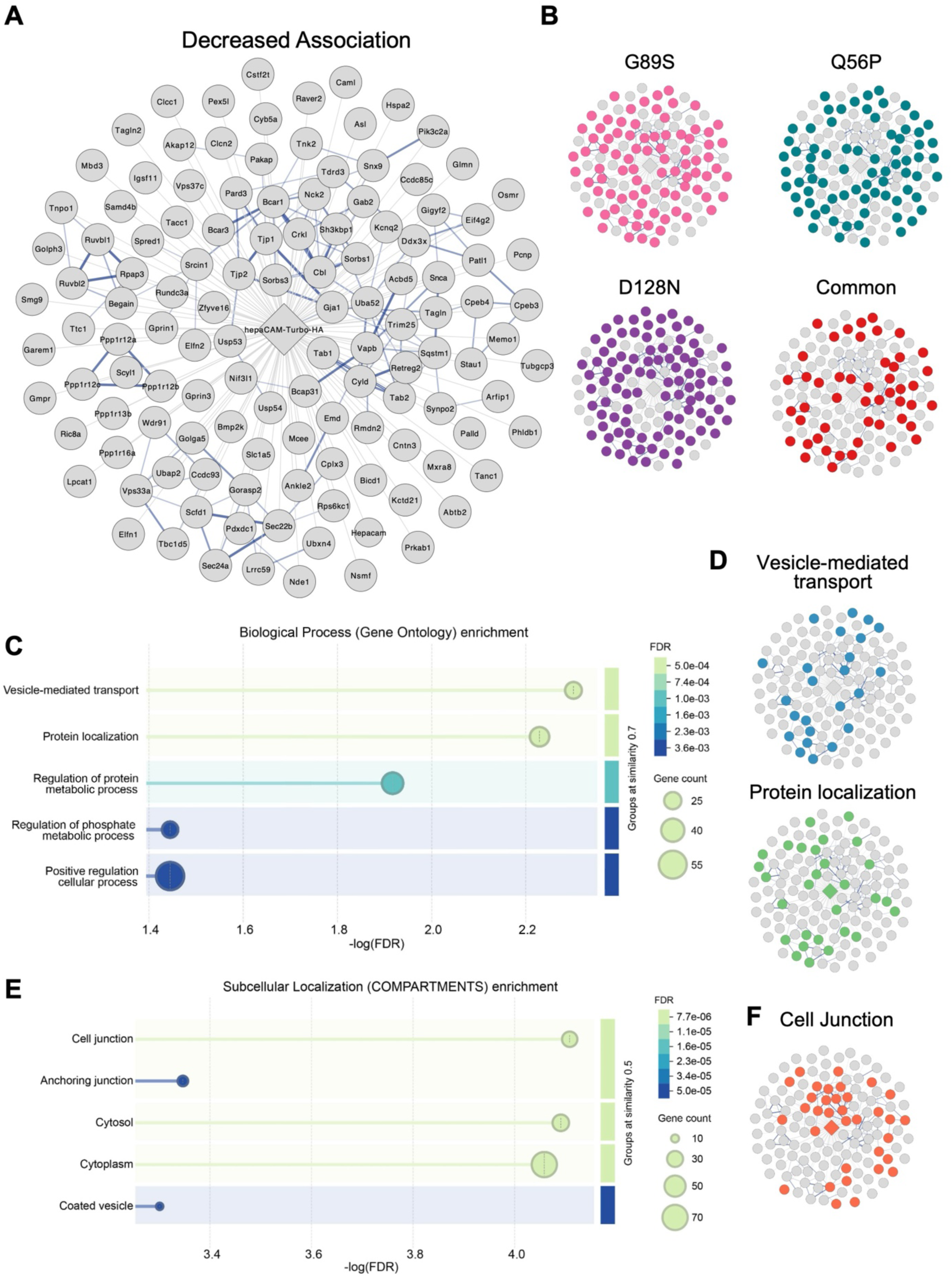
Network analysis reveals hepaCAM mutants decrease interactions with proteins implicated in vesicular transport, protein localization, and cell junctions. A) Significantly decreased proteins from the three mutant conditions vs. WT depicted as an interaction network with hepaCAM-Turbo-HA as the central node. Blue lines represent STRING analysis to detect predicted protein-protein interactions. B) Location of individual mutant vs. WT proteins within the network and proteins common to all three mutant vs. WT networks. C) Gene Ontology analysis of Biological Process enrichment based on -log(FDR). D) Location of proteins associated with top 2 GO terms within the network. E) Gene Ontology analysis of Subcellular Localization (COMPARTMENTS) enrichment based on -log(FDR). F) Location of proteins associated with the top GO term within the network.

For analysis of decreased proteins, we followed the same workflow as above, including all proteins that were significantly increased in at least one mutant condition and performing STRING analysis (**Figure 8A-B**). This network also contained significantly more interactions than expected by chance (PPI p-value = 7.77e-16). The top two biological process GO terms were vesicle-mediated transport and protein localization (**Figure 8C-D**), suggesting that these mutations impair the ability of hepaCAM to associate with protein machinery involved in protein transport and localization. This is consistent with hepaCAM’s role as a chaperone of other transmembrane proteins^15,16^ and the requirement of hepaCAM for proper localization of a growing list of transmembrane proteins^6,11^. Of interest, the top enriched term for subcellular localization was cell junction (FDR = 7.79e-5) (**Figure 8E-F**), which could indicate impaired association between hepaCAM and other cell junction proteins via reduced chaperone function and/or the inability of mutant hepaCAM to localize to cell junctions. The full list of all enriched GO terms is available in **Table S3**. Collectively, this dataset reveals significant changes in association between mutant hepaCAM and numerous transmembrane proteins of physiological relevance, and hints at potentially broad defects in hepaCAM-mediated membrane protein organization.

## Discussion

Here we used new viral tools, super resolution microscopy, and proximity-based quantitative proteomics to examine the impact of three different dominant MLC-causing mutations on astrocytic hepaCAM protein dynamics during brain development. Across all mutants, we found altered subcellular hepaCAM distribution and significant changes in protein interactome. These findings advance our knowledge of hepaCAM protein function during astrocyte development with important implications for understanding how hepaCAM mutants contribute to MLC pathogenesis.

A key finding of our study is that MLC-causing mutations impair hepaCAM’s ability to associate with several key transmembrane proteins and signaling molecules, which may impair hepaCAM trafficking to and function at cell-cell junctions. We found decreased association with known interaction partners, such as CLC2 and Cx43, supporting findings from previous studies and indicating the robustness of our approach. Additionally, we found decreased association with several proteins that are exciting targets for further exploration into the molecular mechanisms of MLC pathogenesis. KCNQ2 is of particular interested, given the causative role of KCNQ2 mutations in DEE, and the prevalence of seizures and signs of ion dysregulation in MLC. Chloride channel CLIC-like protein 1 (CLCC1) was also one of the top decreased proteins in all three mutant conditions. CLCC1 is localized to the ER membrane and is involved in ER calcium ion homeostasis^38,39^. Given hepaCAM’s role at the plasma membrane in regulating ion channel localization and ion homeostasis, investigating the function of hepaCAM and the impact of MLC-causing mutations in the ER membrane is an exciting area for future study. Though we did not find a decreased association with MLC1 in our dataset, this does not preclude altered MLC1 localization or impaired function. The altered distribution of hepaCAM signal in the MLC mutant conditions may substantially impact MLC1 localization and protein dynamics, thereby impacting its function.

The organization of endogenous hepaCAM into discrete puncta throughout the astrocyte membrane and “cupping” interactions with other membrane-targeted proteins (CD44, KCNQ2, Cx43) underscores its emerging role as a molecular organizer of transmembrane proteins. The upstream mechanisms that regulate the expression and localization of hepaCAM itself remain unknown, though our proteomic results provide some insight. Because we chose to compare hepaCAM-TurboID to a cytosolic TurboID control, rather than a membrane-targeted TurboID, we expect to detect enrichment of proteins and GO terms related to membrane protein sorting and transport. Indeed, KEGG pathways analysis of the hepaCAM vs. Turbo interactomes finds significant enrichment of “SNARE interactions in vesicular transport” and “Endocytosis.” (**Table S4**) Interestingly, proteins in these pathways are significantly reduced in abundance in the mutant conditions, which could suggest altered sorting and or trafficking of hepaCAM protein, and could explain the altered subcellular distribution of mutant hepaCAM protein that we observed in our study. In terms of downstream signaling, the hepaCAM C-terminus reveals little in terms of signaling domains and motifs. Previous *in vitro* studies have suggested association between hepaCAM and the actin cytoskeleton^40^, which is of interest given the role of hepaCAM in astrocyte morphogenesis. Our findings point to microtubule dynamics and RhoGTPase signaling as additional downstream pathways of interest. Components of these pathways are significantly reduced in all three MLC mutant conditions (Rmnd2 and Tubgcp3, for example), and Rho GTPases was the top GO term for Reactome pathway in the decreased proteome dataset (FDR = 0.00092, **Table S3**).

There are limitations to our study, which should be considered when interpreting the data. For one, we used viral tools to exogenously express WT and mutant hepaCAM in astrocytes. This approach is advantageous for TurboID, more efficient than generating multiple new mouse lines, and sufficient for producing novel insight into hepaCAM function, protein interaction, and MLC, as we have shown in this study. While exogenous WT hepaCAM expression recapitulated endogenous hepaCAM expression and association with Cx43, it did result in higher overall hepaCAM expression in transduced cells. We compared all mutant conditions to overexpressed WT to control for any caveats associated with this overexpression. Still, there may be unknown consequences of overexpression, and further validation experiments to validate these findings using physiological levels of protein expression could address these concerns. We also acknowledge that while TurboID experiments can approximate and “interactome,” the presence of a protein in a TurboID dataset is not proof of interaction. For interesting candidates, we followed up with STED microscopy of endogenous proteins, where we achieve ∼50 nm resolution, comparable to the resolution of the proximity ligation assay^41,42^. While this does not directly prove interaction, it serves as a valuable *in vivo* validation assay which mitigates caveats of alternative techniques, such as co-immunoprecipitation, wherein direct interactions may not be detected due to their transient nature or due to disruption by the harsh detergent and cell lysis conditions.

A final limitation to consider is that we examined hepaCAM protein function and MLC mutations solely in protoplasmic gray matter astrocytes of the mouse cortex. We chose this region due to both the effectiveness of our tools in targeting and isolating information from this region and our experience studying hepaCAM protein function in developing cortical astrocytes. However, shockingly little is known about astrocytes and astrocyte development in the white matter. This is partially due to the low volume of white matter in the rodent brain (∼10%, compared to 50% in human^43^) and partially due to a lack of tools tailored to targeting and studying white matter astrocytes. Because MLC is a leukodystrophy with significant white matter pathology, an understanding of astrocyte development, hepaCAM protein function, and MLC pathology in white matter is an exciting future avenue of study with significant clinical relevance. Adapting available tools and techniques to target white matter astrocytes is therefore necessary to yield significant new insights into the cellular and molecular drivers of white matter dysfunction in MLC.

## Materials and Methods

### Animals

All mice were used in accordance with the Institutional Animal Care and Use Committee (IACUC) and the UNC Department of Comparative Medicine (IACUC Protocol Numbers 21-116.0 and 24-005.0). Mice were housed in standard conditions with 12-hour day/night cycles. Timed-pregnant CD1 females were obtained from Charles River (RRFD:IMSR_CRL:022). Aldh1l1-GFP transgenic mice were obtained from MMRRC (RRID:MMRRC_011015-UCD). Mice were used for experiments at P21, or as specified in the text and figure legends. For all experiments, mice of both sexes were included in analysis, and we did not observe any influence or association of sex on the experimental outcomes. Criteria for inclusion, exclusion, and randomization is listed for each experiment in specific subsections of the Methods Details section.

### Plasmids

The QuikChange Lightning Site-Directed Mutagenesis Kit (Agilent Technologies 210518) was used to introduce Q56P and D128N mutations into pcDNA3.1-hepaCAM-HA^6^ using the following primers:

Q56P Forward: GCTCTGCTTTCTGTGCCGTACAGCAGTACCAGCA

Q56P Reverse: TACTGCTGTACGGCACAGAAAGCAGAGCCGACTT

D128N Forward: ATCTCCATCACCAACGACACCTTCACTGGGGAGAA

D128N Reverse: CAGTGAAGGTGTCGTTGGTGATGGAGATCTCGAC

For mutagenesis, 10 ng of plasmid DNA was amplified using the mutagenesis primers, reactions were digested with DpnI and transformed into XL10-Gold Ultracompetent Cells on LB-Amp plates. Colonies were isolated and then grown overnight at 37°C in 14 mL round bottom tubes containing 3 mL of Luria-Bertani (LB) media and 300 µg of ampicillin. The ZymoPure II Plasmid Maxiprep Kit (Zymo Research D4202) was used to isolate DNA, and mutations were confirmed by whole plasmid sequencing (Plasmidsaurus). pcDNA3.1-hepacam-HA and pcDNA3.1-hepacam-G89S-HA were generated previously^6^. WT and mutant hepaCAM-HA constructs were subcloned into pZac2.1-gfaABC1D-TurboID-HA^44^ using EcoRI and Not1 restriction enzymes. Final plasmids were sequenced using whole plasmid sequencing (Plasmidsaurus). pZac2.1-gfaABC1D-mCherryCAAX was generated previously^45^.

### Cell Culture

#### Cortical Neuron

Purified rat cortical neurons were prepared as described previously^6^. Briefly, cortices were isolated from P1 rat pups of both sexes, digested in papain (7.5 units/mL), and triturated in low and high ovomucoid solutions. Neurons were resuspended in panning buffer (DPBS (Life Technologies 14287) supplemented with BSA and insulin) and passed through a 20 µm mesh filter (Elko Filtering 03-20/14). Filtered cells were incubated on negative panning dishes coated with Bandeiraea Simplicifolia Lectin 1, followed by goat anti-mouse IgG+IgM (H+L) (Jackson ImmunoResearch 115-005-044), and goat anti-rat IgG+IgM (H+L) (Jackson ImmunoResearch 112-005-044) antibodies, then incubated on positive panning dishes coated with mouse anti-L1 (ASCS4, Developmental Studies Hybridoma Bank, Univ. Iowa) to bind cortical neurons. Adherent cells were dislodged using a P1000 pipet, pelleted (11 min at 200 xg), and resuspended in serum-free neuron growth media (NGM; Neurobasal, B27 supplement, 2mML-Glutamine, 100U/mL Pen/Strep, 1mM sodium pyruvate, 4.2 µg/mL Forskolin, 50 ng/mL BDNF, and 10 ng/mL CNTF). 100,000 neurons were plated onto 12 mm glass coverslips coated with 10 µg/mL poly-D-lysine (PDL, Sigma P6407) and 2 µg/mL laminin and incubated at 37°C in 10% CO_2_. On day *in-vitro* (DIV) 2, half of the media was replaced with NGM Plus (Neurobasal Plus, B27 Plus, 100 U/mL Pen/Strep, 1 mM sodium pyruvate, 4.2 μg/mL Forskolin, 50 ng/mL, BDNF, and 10 ng/mL CNTF) and AraC (10 µM) was added to stop the growth of proliferating contaminating cells. On DIV 3, the media was replaced with NGM Plus. Neurons were fed on DIV 6 and DIV 9 by replacing half of the media with NGM Plus.

#### Cortical Astrocyte

Rat cortical astrocytes were prepared as described previously^6^. P1 rat cortices from both sexes were micro-dissected, digested in papain, triturated in low and high ovomucoid solutions, filtered, and resuspended in astrocyte growth media (AGM; DMEM (Life Technologies 11960), 10% FBS, 10 µM hydrocortisone, 100 U/mL Pen/Strep, 2 mM L-Glutamine, 5 µg/mL Insulin, 1 mM Na Pyruvate, 5 µg/mL N-Acetyl-L-cysteine). Between 15-20 million cells were plated on 75 mm^2^ flasks (non-ventilated cap) coated with poly-D-lysine and incubated at 37°C in 10% CO_2_. On DIV 3, non-astrocyte cells were removed by forceful shaking of closed flasks. Fibroblast elimination was performed by adding AraC to the media on DIV 5. On DIV 7, astrocytes were trypsinized (0.05% Trypsin-EDTA) and plated into 6-well (400,000 cells/well) plates. On DIV 8, cultured rat astrocytes were transfected with shRNA plasmids using Lipofectamine LTX with Plus Reagent (Thermo Scientific) per the manufacturer’s protocol. Briefly, 2 µg total DNA was diluted in Opti-MEM containing Plus Reagent, mixed with Opti-MEM containing LTX (1:2 DNA to LTX) and incubated for 30 minutes at room temperature. The transfection solution was added to astrocyte cultures and incubated at 37C for 3 hours, then replaced with AGM. On DIV 10, astrocytes were trypsinized, resuspended in NGM Plus, plated (20,000 cells per well) onto DIV 10 neurons, and co-cultured for 72 hours.

### Immunocytochemistry

Astrocyte-neuron co-cultures were fixed and stained as described previously^6^. DIV 13 co-cultures were incubated with warm 4% PFA for 7 minutes, washed 3 times with 1x Phosphate Buffer Saline (PBS), and blocked in PBS containing 50% normal goat serum (NGS) and 0.4% Triton X-100 for 30 minutes at room temperature. Samples were washed once more in PBS and incubated overnight at 4°C in chicken anti-GFP (Aves GFP1020, 1:1000) and rat anti-HA (Sigma, 11867423001, 1:500) diluted in antibody blocking buffer (ABB: pH 7.4, 150 mM NaCl, 50 mM Tris, 1% BSA, 100 mM L-lysine, 0.04% sodium azide) containing 10% NGS. The following day, samples were washed 3x with PBS, incubated with goat anti-Chicken 488 and goat anti-rat 594 (Life Technologies, 1:500) diluted in ABB with 10% NGS for 2 hours at room temperature, followed by 10-minute incubation in DAPI (1:50,000) in PBS. Coverslips were washed 3x with PBS and mounted onto glass slides using a homemade glycerol mounting media (20 mM Tris pH 8.0, 90% Glycerol, 0.5% N-propyl gallate). Healthy astrocytes with strong expression of GFP and HA, a single nucleus, and minimal overlap with other GFP+ astrocytes, were imaged at 40x magnification in green, red, and DAPI channels using a Nikon Ti2 widefield fluorescent microscope. Astrocytes containing multiple nuclei, weak GFP expression, or in areas of dense GFP labeled cells were not imaged. The individual acquiring the images was always blinded to the experimental condition. We used ImageJ to calculate the area of GFP and HA signal. The threshold for the GFP channel was adjusted to capture the entire territory of the GFP+ cell. GFP signal from neighboring cells or background was excluded to produce an isolated GFP cell. The area of the GFP+ cell was measured and a mask of the GFP+ cell was created, saved as an ROI, and applied to the HA channel. HA signal outside of the GFP mask was removed, and signal inside of the GFP mask was thresholded to capture HA signal. The total area of HA signal within the GFP mask was calculated using “Analyze Particles” and normalized to the GFP area for each cell. Data were analyzed in Graphpad Prism 10 using a one-way ANOVA with Tukey’s HSD.

### AAV production and administration

The pZac2.1-gfaACB1D plasmids containing TurboID-HA, hepaCAM-TurboID-HA, G89S-TurboID-HA, Q56P-TurboID-HA, D128N-TurboID-HA, or mCherryCAAX were packaged into AAV2/5/PHP.eB capsids by the UNC BRAIN Initiative Viral Vector Core. Purified AAVs were exchanged into storage buffer containing 1 x phosphate-buffered saline (PBS), 5% D-Sorbitol, and 350 mM NaCl. Virus titers (GC/ml) were determined by qPCR targeting the AAV inverted terminal repeats. All viruses were adjusted to the same titer (3.4x10^13^ GC/mL) and 1 µL of virus was injected bilaterally into the cortex of postnatal day 1 (P1) CD1 mice using a Hamilton syringe with a custom removeable needle.

### Immunohistochemistry

#### Sample preparation

For immunohistochemistry, mice were anesthetized with 0.8 mg/kg tribromoethanol (avertin) and perfused with 1x Tris Buffered Saline (TBS)/Heparin followed by ice cold 4% PFA in TBS. Brains were collected and post-fixed overnight in 4% PFA. The following day, brains were rinsed twice with TBS, cryoprotected in 30% sucrose in TBS for 2-3 days, frozen in a medium containing 2:1 30% sucrose to O.C.T. (VWR), and stored at -80 °C. Coronal sections (40 µm thick) were collected using a CryoStar NX50 Cryostat (Thermo Fisher Scientific) and stored at - 25°C in 50% glycerol in TBS. Immunolabeling was performed as described previously^6^. Briefly, slices were washed 3x 10 minutes with TBST (1x TBS containing 0.2% Triton), blocked in blocking solution (TBST containing 10% goat serum), and incubated for two nights in primary antibodies diluted in blocking solution at 4°C while shaking at 100 rpm. The following primary antibodies were used: chicken GFP (Aves GFP1020, 1:1000), rat HA (Sigma 11867423001, 1:500), mouse IgG1 anti-hepaCAM (R&D Systems MAB4108, Confocal 1:500, STED 1:250), rabbit Connexin 43 (Cell Signaling 3512, 1:500), rabbit CD44 (Life Technologies 15675-1-AP, 1:500), guinea pig RFP (Synaptic Systems 390-004, 1:1000), rabbit HA (Cell Signaling C29F4, 1:1000), and rabbit KCNQ2 (Life Technologies PA1929, 1:500). Following primary antibody incubation, sections were washed in 3 x 10 minutes in TBST, incubated in secondary antibody solution (see below for details) for 2-3 hours at room temperature, washed 3 x 10 minutes in TBST, and mounted onto glass slides with homemade mounting medium (20 mM Tris pH 8.0, 90% Glycerol, 0.5% N-propyl gallate) and no. 1.5 coverslips. Coverslips were sealed with nail polish and dried at room temperature before storage at 4°C.

For confocal microscopy, species-specific Alexa-fluor conjugated secondary antibodies produced in goat (Life Technologies) were diluted 1:200 in blocking solution. Isotype-specific secondary antibodies were used for mouse monoclonal primary antibodies (e.g., goat anti-mouse IgG1) to prevent excessive background staining. To detect biotin labeling, fluorophore-conjugated streptavidin (Life Technologies, 1:200) was added to the secondary antibody solution. For three color-STED, the following secondary antibodies were diluted 1:100 in blocking solution: Goat anti-Chicken Alexa-fluor 594 (Life Technologies), Goat anti-Mouse IgG1 Atto 647N (Rockland), and Goat anti-Rabbit CF680R (Biotium).

#### Confocal Image Acquisition and Analysis

For analysis of HA puncta, co-localization of HA and Cx43 puncta, and HA and streptavidin co-co-labeling, z-stack confocal images were acquired on a Leica SPX8 with a 63x objective (15 stacks, 0.34 µm step size). For co-localization of HA and KCNQ2 puncta, z-stack confocal images were acquired on an Olympus Fluoview 3000 with a 60x objective (15 stacks, 0.34 µm step size). Co-localized HA and Cx43 puncta and HA and KCNQ2 puncta were quantified using Synbot^29^ with the following parameters: 2-channel, noise reduction, custom ROI, manual thresholding, minimum pixel = 3, pixel overlap. Prior to running Synbot, the ImageJ tracing tool was used to select the HA-expressing cell(s) in each image, measure the cell area, and save the ROI for use as a custom ROI in Synbot. For each animal, 3 z-stack images (15 slices) were acquired and each image converted in 5 separate maximum projection images (MPI) of 3 slices each for a total of 15 MPIs per animal. For bar graphs, data were analyzed in Graphpad Prism 10 using a one-way ANOVA followed by Tukey’s HSD. Analysis of cumulative frequency distribution used a Kruskal-Wallis test followed by Dunn’s post-test. The individual acquiring the images and performing the analysis was always blinded to the experimental condition. No data were excluded from the analysis.

#### STED Image Acquisition

Three-channel STED images were acquired on the Leica Stellaris 8 FALCON STED using a 100x objective with 2x zoom. Channels were acquired in frame sequential mode to minimize crosstalk. The white light laser allowed for precise selection of excitation wavelength. Software-recommended excitation wavelengths were used for Alexa Fluor 594 and Atto 647N. The excitation wavelength for CF680R was adjusted to 685 to minimized crosstalk with 647N^46^. A 2D-STED donut was applied, and the 775 nm STED depletion laser was set at 90%. Z-stack images were acquired using system-optimized settings for resolution (CD44: 3984x3984; KCNQ2: 4104x4104; image dimension of 58.14x58.14 microns) and z-stack (7 steps, 0.18 µm). Raw images were exported directly to Huygens Essential for deconvolution.

### Brain lysis and streptavidin pulldown

AAV-injected mice were administered subcutaneous biotin (24 mg/kg in PBS) for 3 consecutive days beginning at P18. At P21, mice were anesthetized with avertin, brains were rapidly dissected, and corticies were isolated and flash frozen for storage at -80°C. Brains were collected in batches based on litter, with each litter containing at least 2 brains per condition. Mice were assigned randomly to for each condition. For the proteomics experiment, all 20 streptavidin pulldowns (5 conditions, 4 mice per condition) were performed on the same day in two waves of ten brains, with each wave containing 2 brains per sample (one male and one female). Brain samples were homogenized in a lysis buffer without detergent (50 mM Tris/HCl pH 7.5, 150 mM NaCl, 1 mM EDTA, and 50x protease inhibitors) using a dounce homogenizer and ceramic plunger. An equal volume of 2x RIPA buffer was added (50 mM Tris/HCl pH7.5, 150 mM NaCl, 1 mM EDTA, 0.4% SDS, 2% TritonX100, 2% deoxycholate) to lyse the homogenate and samples were sonicated (3 x 10 second pulses) to further breakdown the tissue. Lysates were centrifugated at 15,000 rpm for 30 minutes at 4°C, and supernatant collected. Sodium dodecyl sulfate (SDS) was added to a final concentration of 1% SDS and samples were heated at 45°C for 45 minutes, then centrifugated at 15,000 rpm for 30 minutes at 4°C. Dynabeads™ MyOne™ Streptavidin T1 (Thermo Fisher 65602, 250 µL slurry per sample) were washed three times in RIPA-IP buffer (50 mM Tris/HCl pH 7.5, 150 mM NaCl, 1 mM EDTA, 2M Urea, 1% NP-40, 0.25% deoxycholate) and added to the samples to incubate overnight at 4°C with rotation (15 rpm). The following day, beads were isolated using a magnetic rack and washed three times (10 minutes each, room temperature, rotating at 15 rpm) with RIPA-IP buffer, followed by three washes with 50 mM ammonium bicarbonate. 10% of each sample was transferred to a new tube and eluted with 70 µL of 2x Sample Buffer^47^ supplemented with 5 mM biotin for use in western blot experiments. The remaining 90% of the sample were resuspended in 50 mM ammonium bicarbonate and stored at -80°C for submission to the UNC Metabolomics and Proteomics Core facility.

### Proteomic analysis

#### Sample Preparation for Affinity Purification Mass Spectrometry Analysis (AP-MS)

Immunoprecipitated samples were subjected to on-bead trypsin digestion, as previously described^48^. After the last wash buffer step, 50 µl of 50 mM ammonium bicarbonate (pH 8) containing 1 µg trypsin (Promega) was added to beads overnight at 37°C with shaking. The next day, 500 ng of trypsin was added then incubated for an additional 3 h at 37°C with shaking. Supernatants from pelleted beads were transferred, then beads were washed twice with 50 ul LC/MS grade water. These rinses were combined with original supernatant, then acidified to 2% formic acid. Peptides were desalted with peptide desalting spin columns (Thermo) and dried via vacuum centrifugation. Peptide samples were stored at -80°C until further analysis.

#### LC/MS/MS Analysis

Each sample was analyzed by LC-MS/MS using an Easy nLC 1200 coupled to a QExactive HF (Thermo Scientific). Samples were injected onto an IonOpticks Aurora Elite TS C18 column (75 μm id × 15 cm, 1.7 μm particle size) and separated over a 120 min method. The gradient for separation consisted of a step gradient from 5 to 36 to 48% mobile phase B at a 250 nl/min flow rate, where mobile phase A was 0.1% formic acid in water and mobile phase B consisted of 0.1% formic acid in 80% ACN. The QExactive HF was operated in data-dependent mode where the 15 most intense precursors were selected for subsequent HCD fragmentation. Resolution for the precursor scan (m/z 350–1700) was set to 60,000 with a target value of 3 × 10^6^ ions, 100ms inject time. MS/MS scans resolution was set to 15,000 with a target value of 1 × 10^5^ ions, 75ms inject time. The normalized collision energy was set to 27% for HCD, with an isolation window of 1.6 m/z. Peptide match was set to preferred, and precursors with unknown charge or a charge state of 1 and ≥ 8 were excluded.

#### Data Analysis

Raw data were processed using the MaxQuant software suite (version 1.6.15.0) for peptide/protein identification and label-free quantitation^49^. Data were searched against a Uniprot Reviewed Mouse database (downloaded 01/2023, containing 17,137 sequences), appended with the HepaCAM sequences, using the integrated Andromeda search engine. A maximum of two missed tryptic cleavages were allowed. The variable modifications specified were: N-terminal acetylation and oxidation of Met. Label-free quantitation (LFQ) was enabled. Results were filtered to 1% FDR at the unique peptide level and grouped into proteins within MaxQuant. Match between runs was enabled. Data filtering and statistical analysis was performed in Perseus software (version 1.6.14.0)^50^. Proteins with log2 fold change (log_2_FC) ≥ 1 in WT or mutant hepaCAM conditions compared to cytosolic TurboID and a p-value < 0.05 were considered statistically and biologically significant. In comparisons of protein abundance between WT and mutant TurboID conditions, proteins with a log_2_FC > 0.585 in the mutant condition compared to WT (corresponding to 50% increase over WT) and p-value < 0.05 were considered significantly increased in abundance. Proteins with a log_2_FC < -0.585 and p-value < 0.05 were considered significantly decreased in abundance. For all comparisons, we excluded proteins that were not significantly enriched above TurboID in at least one condition.

#### Network Analysis

Protein networks of significantly increased and decreased proteins were visualized using Cytoscape (v3.10.3). For each network, hepaCAM-Turbo-HA was set at the source node and significantly increased or decreased proteins were set as the target nodes. The particular mutant conditions in which each target node was significantly changed was input as a target node attribute. STRING analysis was performed on networks using the STRING app within Cytoscape to STRINGify network (confidence score = 0.4), followed by Markov clustering (MCL) using the clusterMaker app (inflation = 3). Following clustering, a perfuse forced directed layout was applied to visually organize the network. Gene Ontology (GO) analysis was performed using the STRING Enrichment App in Cytoscape and repeated on the STRING website to generate graphs of the top 5 GO terms, prioritized by FDR.

### Western blot

Brain lysates or eluted pulldown samples were mixed with 4x Laemmli Sample Buffer (Bio-Rad) containing 5% β-ME and incubated for 45 minutes at 45°C for denaturation. 20 µg of protein or 10 µL of eluate was loaded into 4%-15% gradient pre-cast gels (Bio-Rad) and run at 50V for 5 minutes followed by 150 V for 1 hour. Proteins were transferred to PVDF membrane (Millipore) at 100 V for 1 hour and blocked in Intercept Blocking Buffer (LI-COR). For HA and tubulin immunoblotting of brain lysates, membranes were incubated overnight in primary antibodies (rat HA 1:1000, rabbit tubulin 1:1000) diluted in 3% BSA in TBS-Tween (1x TBS containing 0.1% Tween-20). The next day, membranes were washed 3x with TBS-Tween, incubated in LI-COR IRDye 680RD conjugated secondary antibodies (1:5000 in Intercept Blocking Buffer) for 2 hours at room temperature, and washed two times with TBS-Tween and once with TBS. For streptavidin blotting of eluted sample, membranes were incubated in LI-COR IRDye 800CW Streptavidin (1:5000 in Intercept Blocking Buffer) overnight at 4°C. The next day, the membrane was washed twice with TBS-Tween and once with TBS. Following the TBS wash, membranes were dried and imaged on a LI-COR Odyssey imaging system.

### Statistical Analysis

All statistical analyses were performed in GraphPad Prism 10, with the exception of proteomics analysis which was performed as described in the proteomic methods subsection. For each experiment, the number of subjects and specific statistical tests are included in the figure legend and data are represented as mean ± standard error of the mean with exact P-values shown. Sample sizes were determined based on previous experience for each experiment and no statistical methods were used to predetermine sample size. Specific details for inclusion, exclusion, and randomization are included in specific method subsections.

