## Supplemental Figures 1-3 for "Dominant MLC-causing mutations alter hepaCAM subcellular localization and protein interactome in astrocytes of the developing mouse cortex"

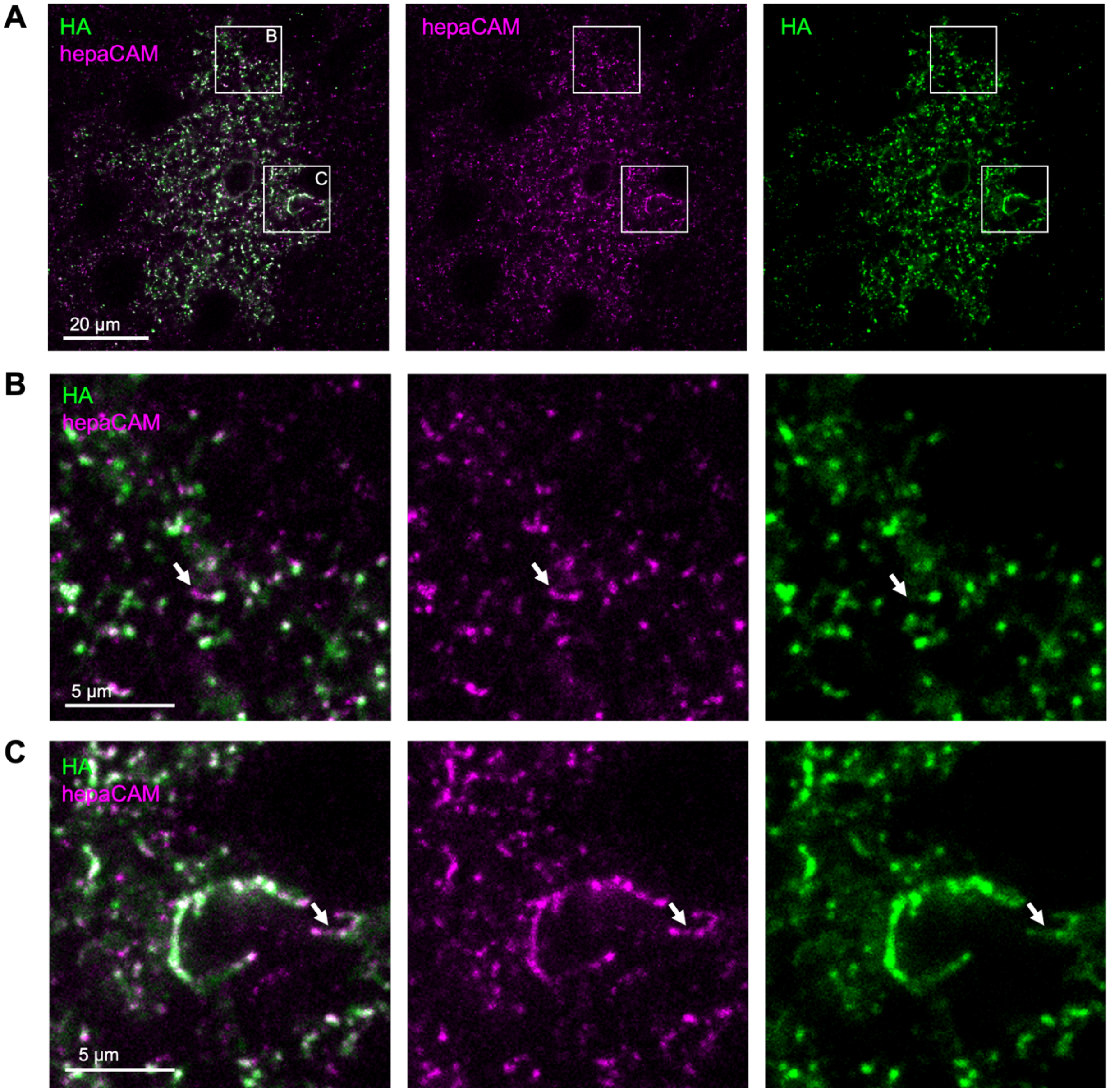


**Supplemental Figure 1: Exogenous hepaCAM-Turbo-HA displays similar subcellular localization to endogenous hepaCAM**

A) Representative maximum-projection images (3-slices, 0.34 μm step size) of transduced astrocytes in layer 5 of the mouse VCX at P21 with HA in green and endogenous hepaCAM in magenta. B) Inset from A depicting localization of hepaCAM and HA within the astrocyte arbor. Arrow denotes an example of magenta signal that does not co-localize with green HA signal. C) Inset from A depicting localization of hepaCAM and HA at the astrocyte endfoot. Arrow denotes an example of magenta signal that does not co-localize with green HA signal.

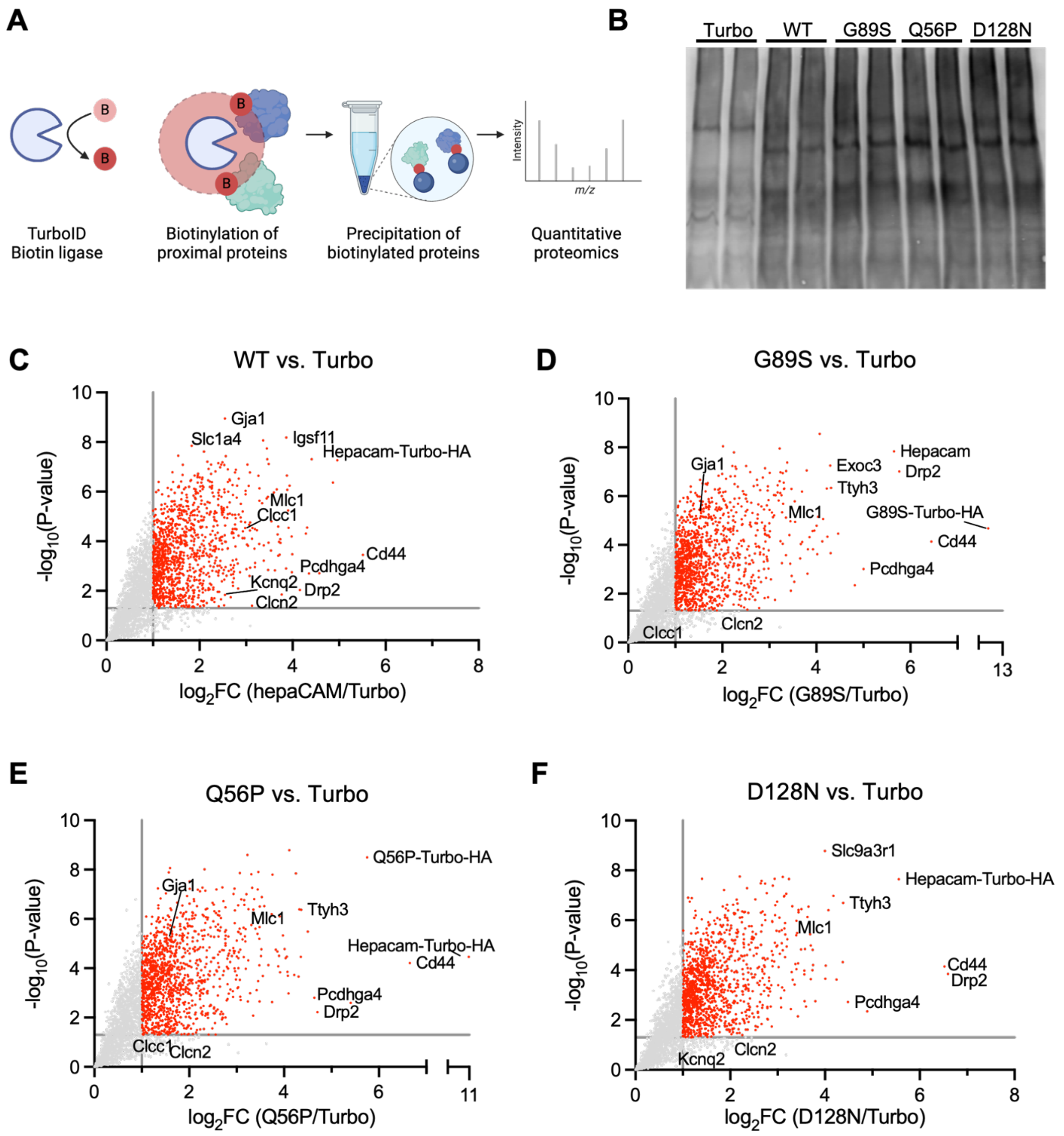


**Supplemental Figure 2: WT and mutant proteomes compared to TurboID control**

A) Schematic of TurboID workflow. B) Western blot of TurboID pulldowns probed with streptavidin 680. C-F) X-Y plots depicting significantly enriched proteins over TurboID control for C) WT, D) G89S, E) Q56P, and F) D128N. Red dots indicate proteins significantly enriched in the hepaCAM-TurboID condition ($p < 0.05$, $log_2FC > 1$).

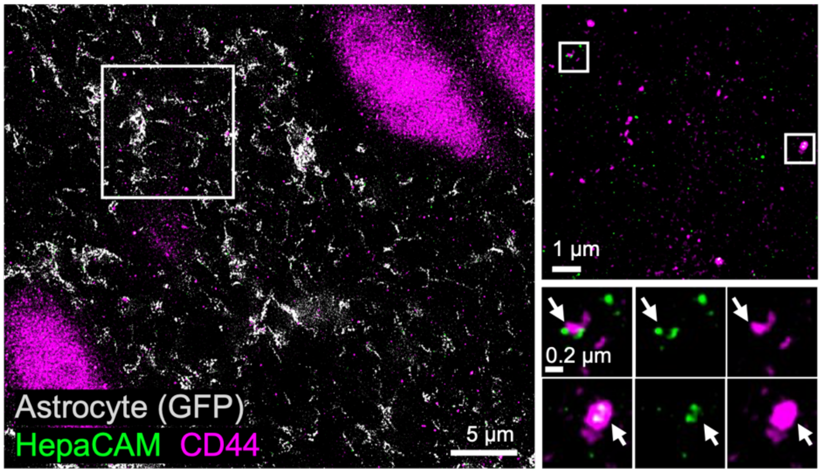


**Supplemental Figure 3: Co-localization of endogenous hepaCAM and CD44 in super resolution**

Representative three-color STED images of astrocytes in L5 VCX from Aldh1L1eGFP mice. GFP-labeled astrocyte (gray), endogenous hepaCAM (green), CD44 (magenta).
